# Expanding the phage galaxy: Isolation and characterization of five novel *Streptomyces* siphoviruses Ankus, Byblos, DekoNeimoidia, Mandalore, and Naboo

**DOI:** 10.1101/2024.02.16.580700

**Authors:** Sebastian H. Erdrich, Tom Luthe, Larissa Kever, Biel Badia Roigé, Borjana Arsova, Eva Davoudi, Julia Frunzke

## Abstract

Key features of the actinobacterial genus *Streptomyces* are multicellular, filamentous growth and production of a broad portfolio of bioactive molecules. These characteristics appear to play an important role in phage-host interactions and are modulated by phages during infection. To accelerate research of such interactions and the investigation of novel immune systems in multicellular bacteria, phage isolation, sequencing, and characterization are needed. This is a prerequisite for establishing systematic collections that appropriately cover phage diversity for comparative analyses. As part of a public outreach programme within the priority programme SPP 2330, involving local schools, we describe the isolation and characterization of five novel *Streptomyces* siphoviruses infecting *S. griseus, S. venezuelae*, and *S. olivaceus*. All isolates are virulent members of two existing genera and, additionally, establish a new genus in the Stanwilliamsviridae family. In addition to an extensive set of tRNAs and proteins involved in phage replication, about 80% of phage genes encode hypothetical proteins, underlining the yet underexplored phage diversity and genomic dark matter still found in bacteriophages infecting actinobacteria. Taken together, phages Ankus, Byblos, DekoNeimoidia, Mandalore, and Naboo expand the phage diversity and contribute to ongoing research in the field of *Streptomyces* phage-host interactions.

## 1. Introduction

*Streptomyces*, belonging to the phylum of Actinobacteria, are of significant interest due to their developmental life cycle as well as their sophisticated specialized metabolism (Barka et al., 2016; Chater, 2016; Shepherdson et al., 2023). Unlike most bacteria dividing by binary fission, *Streptomyces* are characterized by a multicellular development starting with the germination of spores, forming a network of vegetative hyphae. Stressful conditions initiate the transition to a reproductive aerial mycelium, further differentiating into spore chains (Bush et al., 2015; Flärdh and Buttner, 2009).

Production of bioactive compounds is closely linked to the developmental program and is usually initiated upon the developmental switch to the growth of aerial hyphae (Rigali et al., 2008; Yagüe et al., 2012). On average, *Streptomyces* species encode 31 distinct specialized metabolites relevant to medicine, biotechnology, and agriculture (Alam et al., 2022; Nikolaidis et al., 2023; Otani et al., 2022). However, many biosynthetic gene clusters (BGCs) responsible for the synthesis of such bioactive compounds are silent under laboratory conditions, limiting the full appreciation of the chemical potential of *Streptomyces*. Developing ways to activate these BGCs has gained significant interest in recent years, which might allow access to new antimicrobial compounds (Karimian et al., 2024; Liu et al., 2021).

Besides genetic engineering approaches, microbial interactions have the potential to be a potent strategy for stimulating BGCs (Netzker et al., 2018). Surprisingly, phages, as the most abundant bacterial predator, have not yet been considered as a potential trigger of secondary metabolism. However, recent studies show the antiphage properties of *Streptomyces*-derived aminoglycosides and anthracyclines (Jiang et al., 2020; Kever et al., 2022; Kronheim et al., 2018), as well as initial phenomenological and transcriptional clues indicating that phage infection actually triggers the production of specialized metabolites (Hardy et al., 2020; Kronheim et al., 2023; Luthe et al., 2023).

Although phages infecting *Streptomyces* are still underexplored, efforts to isolate and characterize such actinobacteriophages have accelerated significantly, mainly initiated by the Science Education Alliance-Phage Hunters Advancing Genomics and Evolutionary Science (SEA-PHAGES) program in the US (Hatfull, 2015). While the resulting Actinobacteriophage Database (phagesdb.org, February 2024) currently holds information about 4850 sequenced bacteriophages infecting 13 different Actinobacteria genera, there is considerable bias towards those infecting *Mycobacterium* species (Hatfull, 2020). Following the success of this science program, we here report the characterization and genome analysis of five novel *Streptomyces* phages isolated as part of the high school outreach program “Going Viral”. This program was organized in the frame of the priority program SPP 2330 (www.spp2330.de) together with JuLab at the Forschungszentrum Jülich.

## 2. Material and methods

### 2.1 Bacterial strains and growth conditions

For liquid cultures, all *Streptomyces* strains were inoculated from spore stocks and grown in GYM medium (per litre: 4 g glucose, 4 g yeast extract, 10 g malt extract, pH = 7.3). Unless mentioned otherwise, cultivations were performed overnight at 30 °C and 170 rpm. Growth on plates was performed using GYM agar (adding 1.5 % agar-agar and 20 mM CaCO_3_). For infections, GYM soft agar (0.4 % agar-agar) was mixed with the host strain and used as top agar in double agar overlays.

### 2.2 Phage isolation and propagation

For phage isolation, *S. griseus* DSM 40236, *S. venezuelae* NRRL B-65442, and *S. olivacues* DSM 41536 were used as host strains. The bacteriophages were isolated by adding SM buffer (50 mM Tris-HCl, 100 mM NaCl, 8 mM MgSO_4_, pH = 7.5) to soil samples collected within a radius of 20 km around the Forschungszentrum Jülich. After incubation for three hours on a rock shaker, the samples were centrifuged at 4,000 x g for 20 minutes to remove solid particles. The supernatants were filtered through 0.2 µm pore size membrane filters (Sarstedt; Filtropur S, PES). For phage enrichment, filtered supernatant solution was added to 5x concentrated GYM medium with 500 µl of a *Streptomyces* overnight culture, resulting in a 1x GYM concentration. This enrichment culture was incubated at 30 °C and 170 rpm overnight. Afterwards, the culture was centrifuged at 4,000 x g for 20 min to collect the supernatant, which was subsequently filtered with 0.2 µm filters. Serial dilutions of the enriched supernatant were made in SM buffer and spotted on double agar overlay plates containing the host bacterium in the top agar (OD_450_ = 0.4). Plaques were visible after overnight incubation. Phage samples were purified by restreaking single plaques at least three times. Plaque morphology had to stay constant before a sample was considered a single phage isolate (Kauffman and Polz, 2018). Harvesting of purified phage particles was performed after overnight incubation. The top agar was solubilized by adding 5 mL SM buffer and two hours incubation at RT on a rock shaker. The solution was subsequently transferred into a falcon tube and centrifuged at 4,000 g for 20 min to remove the residual amounts of top agar. The supernatant was filtered through 0.2 µm filters and stored at 4 °C. A dilution series was spotted on overlay agar for titer determination, and the visible plaques at the highest dilution were counted.

### 2.3 Negative staining transmission electron microscopy (TEM) of phage virions

For electron microscopy of single phage particles, 3.5 µL purified phage suspension was absorbed on a glow discharged (15 mA, 30 s) carbon-formvar coated copper grid (CF300-CU, Carbon film 300 mesh copper) and were subsequently washed twice in water, directly stained for 30 sec in 6 µL of 2 % (wt/vol) uranyl acetate and left for drying. The negative-stained samples were examined on a Talos L120C G2 transmission electron microscope (Thermo Fisher Scientific, Dreieich, Germany), which was operated at 120 kV (LaB6 / Denka).

### 2.4 Host range determination

For host range determination and efficiency of plating testing, phages were amplified on their respective isolation host and serially diluted in SM buffer. Then, 2 µL of each dilution were spotted on bacterial lawns prepared as overlays from soft agar containing either *S. griseus* DSM 40236, *S. venezuelae* NRRL B-65442, *S. olivacues* DSM 41536, *S. albidoflavus* M145 (formerly *S. coelicolor* M145*), S. xanthochromogenes* DSM 40111, or *Streptoalloteichus tenebrarius* DSM 40477. Spotting was performed in duplicates. A species was considered part of the host spectrum of the phage if single plaques were visible. The efficiency of plating was calculated relative to the isolation host.

### 2.6 DNA isolation

Phage DNA was isolated by treating 2 mL of phage solution with 1 U/μL DNase I (Invitrogen, Carlsbad, CA, USA) to remove free DNA. The further steps of DNA isolation were performed with the Norgen Biotek Phage DNA Isolation Kit (Norgen Biotek, Thorold, Canada) according to the manufacturers protocol.

### 2.7 DNA sequencing and genome assembly

For genome sequencing, sample quality control, library preparation and sequencing were performed by GENEWIZ, Leipzig, using the Illumina MiSeq platform with a read length of 2 × 150 bp (Illumina). A subset of 100,000 reads was sampled for each phage, and a de novo assembly was performed with CLC genomics workbench 20.0.4 (QIAGEN, Hilden, Germany). Finally, contigs were manually curated and checked for coverage.

### 2.8 Gene prediction and functional annotation

The phage open reading frames (ORFs) were predicted with Pharokka v 1.3.2 (Bouras et al., 2023) in terminase reorientation mode using PHANOTATE (McNair et al., 2019), tRNAs were predicted with tRNAscan-SE 2.0 (Chan et al., 2021), tmRNAs were predicted with Aragorn (Laslett, 2004), and CRISPRs were checked with CRT (Bland et al., 2007). Functional annotation was generated by matching each CDS to the PHROGs (Terzian et al., 2021), VFDB (Chen, 2004) and CARD(Alcock et al., 2019) databases using MMseqs2 (Steinegger and Söding, 2017) PyHMMER (Larralde and Zeller, 2023). Contigs were matched to their closest hit in the INPHARED database (Cook et al., 2021) using mash (Ondov et al., 2016). Plots were created with the pyCirclizen package. Genome termini classes were determined using Phage Term (Garneau et al., 2017), and parameters were set by default. Phage lifestyle was predicted by the machine learning based program PhageAI (Tynecki et al., 2020) using default parameters and further confirmed by the absence of integrase genes inside the genomes.

The annotated genomes were deposited in GenBank under the following accession numbers: PP171438 (Ankus), PP171439 (Byblos), PP171440 (DekoNeimoidia), PP171441 (Mandalore), PP171442 (Naboo).

### 2.9 Genome comparison and classification

The novel phage isolates were classified based on nucleotide sequence comparison against known *Streptomyces* phages (Hardy et al., 2020). Closely related bacteriophages were recovered from the NCBI nucleotide blast searches. The average nucleotide identities (ANI) were calculated by pairwise comparison of our five novel phages to the reference genomes using VIRIDIC (Moraru et al., 2020) with default settings (70 % Genus threshold and 95 % ANI as species threshold). Phylogenetic analysis was performed in MEGA X (Kumar et al., 2018). Alignment was done with MUSCLE on default parameters (Edgar, 2004). The tree was drawn using the WAG+G+F algorithm (Whelan and Goldman, 2001) as a best fit model with 100 bootstrap replicates. The final visualization of the tree was done with iTOL (Letunic and Bork, 2021).

## 3 Results

### 3.1 Phage isolation, morphology and host range

Five novel bacteriophages infecting *Streptomyces* species were isolated from environmental soil samples collected in a 20 km radius around the Forschungszentrum Jülich (Germany) by high school students as part of the citizen science project “Going Viral” in the frame of the DFG-funded priority programme SPP 2330 (“New concepts in prokaryotic virus-host interaction”, www.spp2330.de). Phages Ankus, Byblos, and DekoNeimoidia were isolated on *Streptomyces griseus*, phage Mandalore was obtained with *Streptomyces venezuelae*, and Naboo was isolated on *Streptomyces olivaceus* (Figure 1A). All phages form clear plaques (Figure 1B). The size of 15 randomly measured plaques per phage vary drastically between the phages, even on the same host. After 42 h of incubation, plaques of Ankus and DekoNeimoidia on *S. griseus* are comparable in size, being 1.21 mm^2^ (±0.57 mm^2^) and 1.15 mm^2^ (±0.34 mm^2^), respectively. Byblos forms very large, typically round plaques averaging at 6.48 mm^2^ (±2.24 mm^2^). In contrast, plaques formed by Mandalore after 42 h incubation on *S. venezuelae* are the smallest, with an average size of 0.07 mm^2^ (±0.03 mm^2^). The plaques of Naboo on *S. olivaceus* have an average size of 0.83 mm^2^ (±0.32 mm^2^). TEM analysis of purified phage particles showed that all phages have a typical siphoviral morphology with icosahedral capsids of sizes ranging from 66-81 nm (Supplementary Table S1) and non-contractile tails between 343-359 nm (Figure 1C).

**Figure 1:**
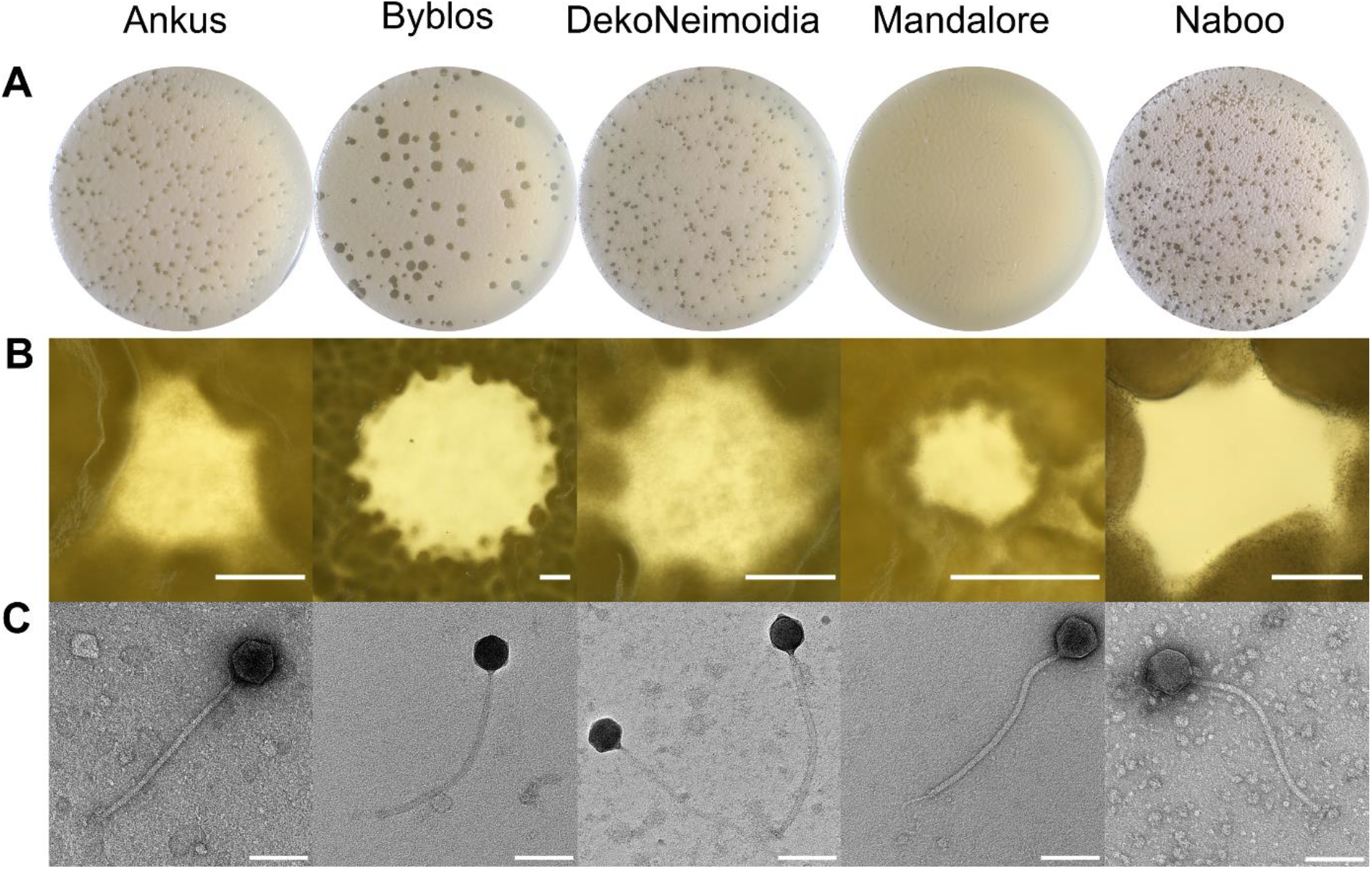
Phage and plaque morphology of novel *Streptomyces* phages. (**A**) Plaque morphologies of the five different phages on double agar overlays. The double agar overlays for phages Ankus, Byblos and DekoNeimoidia were performed with *Streptomyces griseus*; for Mandalore, *Streptomyces venezuelae* was used, and Naboo was plated on *Streptomyces olivaceus*. Images were taken after 42 h of incubation at 30 °C. (**B**) Stereo microscope images of single representative plaques for each phage were taken after 42 h. Scale bar: 500 µm. (**C**) Transmission electron microscopy (TEM) images of virion particles. The phage isolates were negative stained with uranyl acetate. Scale Bar: 100 nm.

While phages are known to be very host-specific and, therefore, often have a narrow host range, some exceptional phages are polyvalent and can infect many strains of the same species. We assessed the host range of our five phages by spotting them on bacterial lawns of five *Streptomyces* species, including the three isolation hosts and one additional *Streptoalloteichus* species (Table 1). Of the isolated phages, Mandalore solely infected its isolation host, *S. venezuelae*. Interestingly, phages isolated on *S. griseus* were able to infect *S. olivaceus* and vice versa. However, Naboo, isolated on *S. olivacues*, showed a higher EOP on *S. griseus*, whereas Ankus, Byblos, and DekoNeimoidia were strongly impaired on their second host. Notably, Ankus, DekoNeimoidia, and Naboo spotting of undiluted phage suspension resulted in a visible morphological change of the *S. albidoflavus* (previously *S. coelicolor)* bacterial lawn but no clear lysis after 42 h. Compared to the other phages and dilutions spotted, here red coloration of the lawn was enhanced. Overall, the host range of the isolated phages appeared to be rather narrow.

**Table 1:**
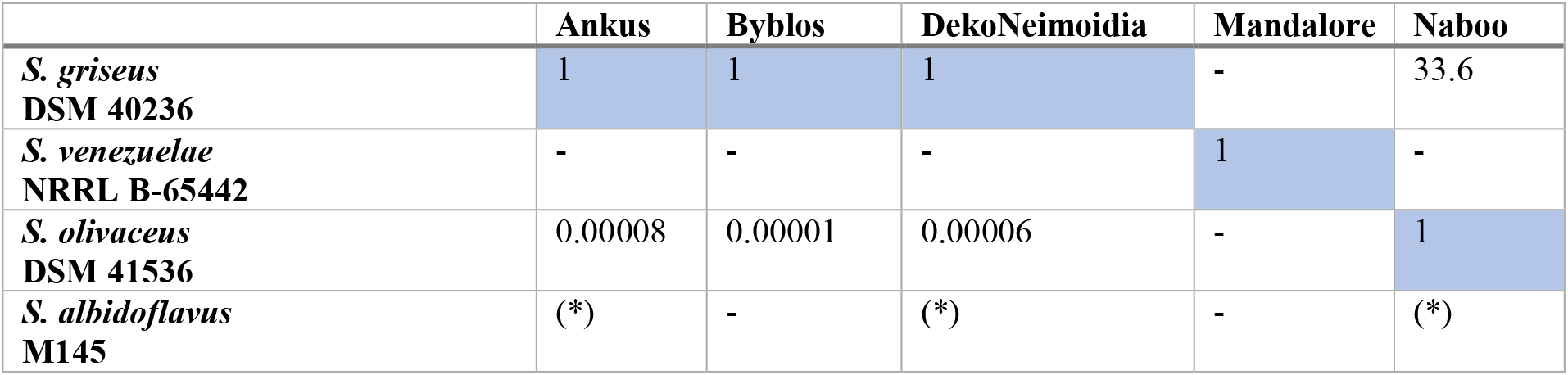
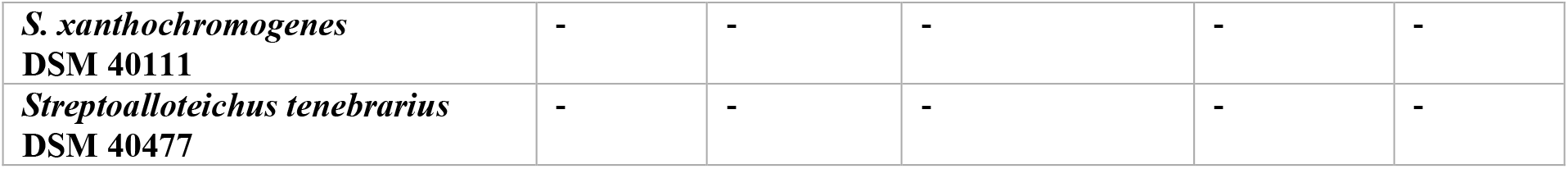
Host range determination. The host range of the five phages was determined by spotting serial dilutions of the phages on lawns of different *Streptomyces* species. EOP was calculated for single plaques in comparison to the isolation host highlighted in blue. (*) indicates no lysis but a morphological reaction of the lawn to the undiluted spot.

|  | Ankus | Byblos | DekoNeimoidia | Mandalore | Naboo |
| --- | --- | --- | --- | --- | --- |
| *S. griseus*<br>DSM 40236 | 1 | 1 | 1 | - | 33.6 |
| *S. venezuelae*<br>NRRL B-65442 | - | - | - | 1 | - |
| *S. olivaceus*<br>DSM 41536 | 0.00008 | 0.00001 | 0.00006 | - | 1 |
| *S. albidoflavus*<br>M145 | (*) | - | (*) | - | (*) |
| *S. xanthochromogenes*<br>DSM 40111 | - | - | - | - | - |
| *Streptoalloteichus tenebrarius*<br>DSM 40477 | - | - | - | - | - |

### 3.2 Genome sequencing and features

The phage isolates were sequenced using Illumina Mi-seq short-read technology. Genomic features of the five isolated phages are summarized in Table 2. Briefly, they have genome sizes ranging from 120 to 127 kb, the GC content varies in the range of 46 to 52%, and each phage genome was predicted to contain 233-252 ORFs. All phages are predicted to be virulent and encode an extensive amount of tRNAs ranging from 32 to 44 genes, spanning each all of the 20 essential amino acids except for Mandalore, which lacks a tRNA for cysteine. Except for Byblos, most of these tRNA genes are located in the first third of the genomes. Additionally, all phages have a set of genes relevant to DNA metabolism, encoding their own DnaB-like replicative helicase, single-strand DNA binding protein, a DNA primase, and DNA polymerase. For Ankus, Byblos, and DekoNeimoidia infecting *S. griseus*, a gene coding for a DNA polymerase exonuclease subunit was found as well. Interestingly, Mandalore and Naboo harbour a gene coding for an FtsK-like protein, and Ankus, Byblos, DekoNeimoidia, and Naboo additionally encode WhiB-like and Lsr2-like transcriptional regulators, which are widespread in actinophages (Sharma et al., 2021). All genomes contain almost 80% of hypothetical proteins, emphasizing the large amount of ‘dark matter’ in actinobacteriophage genomes.

**Table 2:** Basic genomic features of the five novel phages. a) Open reading frames (ORFs) were predicted with Pharokka v 1.3.2 (Bouras et al., 2023), described in more detail in the material and methods section. b) Genome termini classes were determined using PhageTerm (Garneau et al., 2017). c) Phage lifestyle was predicted by the machine-learning-based program PhageAI (Tynecki et al., 2020). The absence of intergrase genes further confirmed the lytic lifestyle.

| Phage Name | Accession Number | Reference Host | Genome Size (bp) | GC Content (%) | ORF Number<sup>a)</sup> | Genome Termini Class<sup>b)</sup> | Lifestyle Prediction<sup>c)</sup> |
| --- | --- | --- | --- | --- | --- | --- | --- |
| Ankus | PP171438 | *S. griseus* | 121,275 | 52 | 236 | DTR (long) | virulent |
| Byblos | PP171439 | *S. griseus* | 127,548 | 52 | 249 | DTR (long) | virulent |
| DekoNeimoidia | PP171440 | *S. griseus* | 127,012 | 52 | 239 | DTR (long) | virulent |
| Mandalore | PP171441 | *S. venezuelae* | 120,241 | 46 | 252 | DTR (long) | virulent |
| Naboo | PP171442 | *S. olivaceus* | 120,050 | 48 | 233 | N.A. | virulent |

The phage genomes show a phage typical modular clustering of functional genetic units, fulfilling similar functions during host takeover and production of phage progeny (Supplementary Figure S1). To compare the five phages on protein coding level simmilarity of the predicted coding sequences, gene products were compared (Figure 2). The phages Ankus, Byblos and DekoNeimoidia show a high conservation on protein level, but especially Byblos features drastic genome rearrangements, swapping the starting and ending halves of its genome compared to Ankus and DekoNeimoidia. Notably, most of the structural proteins (green) share a high level of similarity between all five phages. Mandalore and Naboo appear to share further genes featuring significant similarity throughout their genomes.

**Figure 2:**
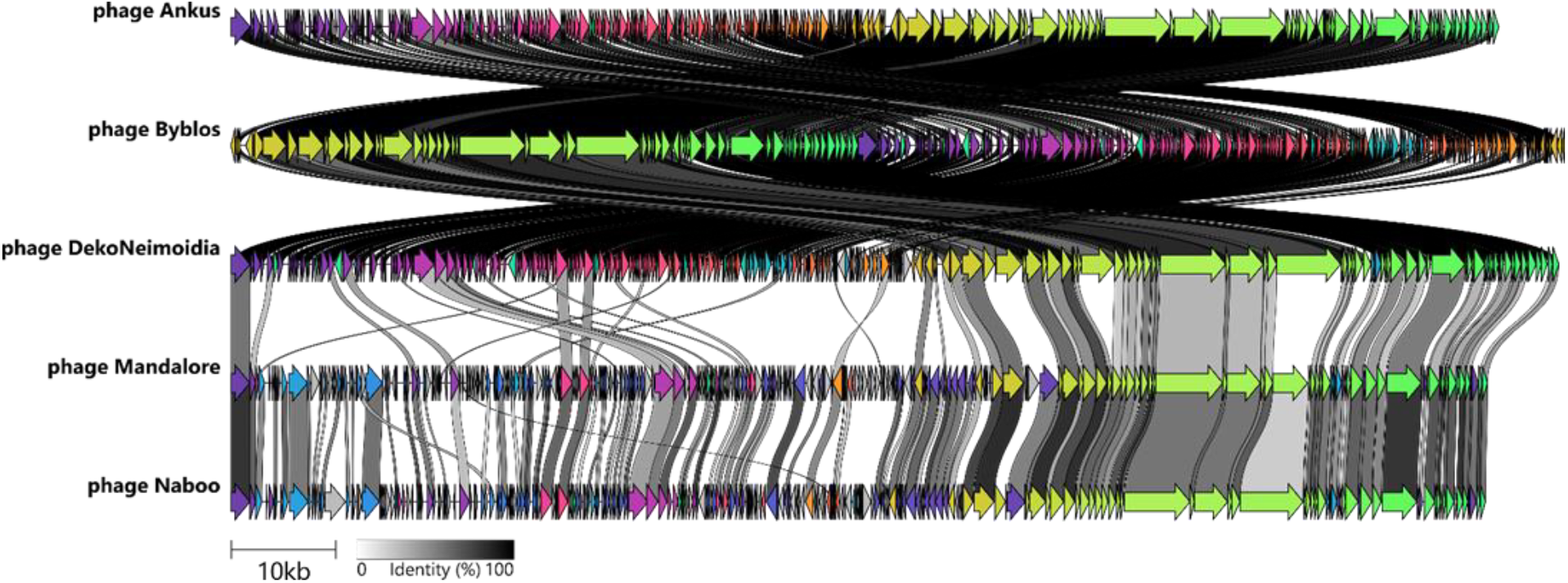
Genome comparison on CDS level. The coding sequences (CDS) of our isolated phages were compared using the clinker pipeline (Gilchrist and Chooi, 2020) to cluster them in groups by similarity (each colour represents one group). The percent identity is indicated in shades of grey. The genomes are represented linear, and the direction of the arrows is in line with the transcription direction of each CDS.

### 3.3 Average nucleotide identity (ANI), phylogenetic analysis and taxonomy

The diversity within our isolates is further investigated by a clustering performed against 11 representative phages infecting *Streptomyces* species, which were obtained from NCBI, based on their genetic relatedness to our five isolates. According to our ANI analysis (Figure 3A), all five isolated phages belong to the family of *Stanwilliamsviridae* within the *Caudoviricetes* class. Herein, Mandalore and Naboo fall into the subfamily of *Boydwoodruffvirinae*, as they are closest related to phage Coruscant (73%, NC_070782) and Tomas (94.1%, NC_070781), respectively, forming the *Coruscantvirus* and *Tomasvirus* genera. Ankus, Byblos, and DekoNeimoidia are closely related, with Ankus sharing an average nucleotide identity of 66.8% with phage Circinus (MK620896). According to the criteria of the ICTV’s Bacterial and Archaeal Viruses Subcommittee for a genus to have at least 70% nucleotide identity between its members (Turner et al., 2021), these three phages thereby potentially create a new genus within the *Loccivirinae* subfamily of *Stanwilliamsviridae*. However, while manual BLAST comparison supports the low level of similarity between Ankus and its closest found relatives, Circinus and BillNye in the *Wilnyevirus* genus (85% identity over 71% query coverage), web-based OrthoANIu algorithm (Yoon et al., 2017) resulted in 81% similarity. Using the helicase protein encoded by all of the 16 phages used here, we additionally performed a phylogenetic analysis (Figure 3B). The maximum-likelihood tree shows a clear distinction between Mandalore and Naboo as well as Ankus, Byblos, and DekoNeimoidia grouped into the two subfamilies of *Stanwilliamsviridae*. Additionally, it supports classification into the aforementioned genera. Furthermore, this approach strengthens the assignment of our novel phages to the known actinobacteriophage clusters as established by The Actinobacteriophage Database (phagesdb.org, Hatfull, 2020). This resulted in the association of Mandalore and Naboo with cluster BE, subcluster BE2, while Ankus, Byblos, and DekoNeimoidia belong to the BK cluster, subcluster BK2.

**Figure 3:**
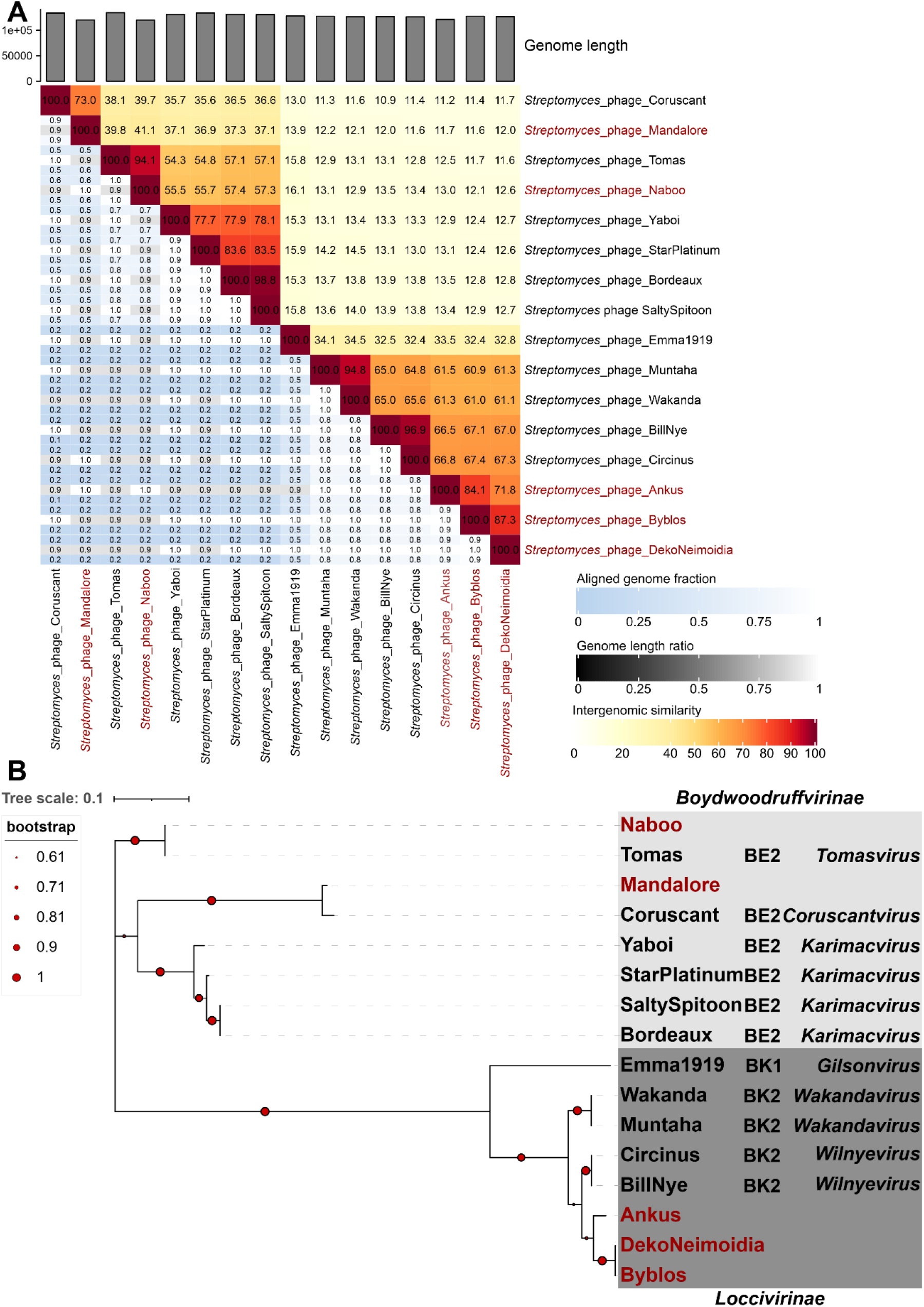
Nucleotide-identity and protein phylogeny enable classification of five novel *Streptomyces* phages. Genomes were acquired from NCBI based on relatedness to our phages. The novel isolated phages are labelled in red. (**A**) The average nucleotide identity (ANI) analysis was performed using VIRIDICT (Moraru et al., 2020). (**B**) Phylogenetic analysis of the phage-encoded helicase proteins. Amino acid sequences were retrieved from the NCBI and aligned using MUSCLE (Edgar, 2004). The best-fit model WAG+G+F (Whelan and Goldman, 2001) was used to draw the shown maximum-likelihood tree with 100 bootstrap replicates indicated as red dots on the branches. Additional annotation provides information on the actinobacteriophage cluster and the associated genera in the two relevant subfamilies. Visualization was done using iTol (Letunic and Bork, 2021). The tree scale represents 0.1 substitutions per site.

## 4. Discussion

This study presents the genomes and basic characteristics of five novel bacteriophages infecting *Streptomyces*. Ankus, Byblos, DekoNeimoidia, Mandalore, and Naboo are all representatives of the largest actinobacteriophage group featuring siphovirus morphology (Hatfull, 2020). They are, in general, larger in size and lower in GC content than the average when compared to the known *Streptomyces* phage diversity (phagesdb.org). Interestingly, this pattern holds true for many *Streptomyces* phages in the Actinobacteriophage Database, where phage genomes larger than 100,000 bp have GC contents below 53%. These phages, like our new isolates, fall into the BE and BK clusters, further supporting our classification (phagesdb.org; Hatfull, 2020). All of them exhibit an overall narrow host range based on the present analysis. Despite systematic phage host range analyses are not commonly standardized or done at all, it appears that *S. venezuelae* phages typically are very specific for their host as other phages like Alderaan and Coruscant (Hardy et al., 2020), which also show the same specificity infecting only one species like our newly isolated phage Mandalore. Interestingly, the observed host range of Ankus, Byblos, DekoNeimoidia and Naboo, all infecting both *S. griseus* and *S. olivaceus*, could be a more general feature. This could hint towards a common receptor present in both cell envelopes, as phage adsorption is the first and major determinant of host range (De Jonge et al., 2019; Magill and Skvortsov, 2023).

Our analyses reveal notable differences between Ankus, Byblos, DekoNeimoidia, and their closest relatives from the *Wilnyevirus* genus. Evaluating the average nucleotide identity and phylogeny of the helicase proteins on the basis of the criteria given by the ICTV’s subcommittee (Turner et al., 2021) with respect to the differences between already established genera (e.g. *Wilnyevirus* and *Wakandavirus*), we propose the formation of a new genus within the *Loccivirinae* subfamily encompassing phages Ankus, Byblos, and DekoNeimoidia.

One major asset of all five phages is the presence of genes relevant for DNA metabolism (i.e. phage replication). This is a common feature of the phages in the *Stanwilliamsviridae* family, which harbour helicases, DNA primases, polymerases, and ssDNA binding proteins essential for forming independent replisomes, as described for different model bacteriophages like T4 or T7 (Benkovic and Spiering, 2017; Magill et al., 2018). However, these proteins have not yet been further investigated in this group of bacteriophages infecting *Streptomyces*. Still, they could potentially provide interesting characteristics in terms of, for example, speed, temperature sensitivity, incorporation of non-canonical nucleotides, or overall replication mechanisms known from diverse phage-encoded polymerases (Morcinek-Orłowska et al., 2022). These features could be harnessed for molecular biology and biotechnological applications.

Furthermore, Ankus, Byblos, DekoNeimoidia, Mandalore, and Naboo encode FtsK-like, Lsr2-like, and WhiB-like proteins known from host regulatory networks in *Streptomyces*, and which are found in many other actinobacteriophages (Bush, 2018; Chen and Banfield, 2024; Hardy et al., 2020; Sharma et al., 2021). Some other genomic features include a MazG-like pyrophosphatase in phage Mandalore, potentially interfering with the host abortive infection mechanism (Gross et al., 2006; Harms et al., 2018) and a Cas4 family exonuclease encoded by Naboo that could lead to host spacer acquisition and subsequently autoimmunity of the bacterial host (Hooton and Connerton, 2015). Remarkably, all phages encode a substantial number of tRNAs, ranging from 32 to 44 genes, covering all 20 essential amino acids. The exception is Mandalore, which lacks a tRNA for cysteine. The extensive repertoire of tRNA genes could be employed to enhance gene expression in hosts with varying codon usage patterns or to counteract potential defense systems based on tRNA degradation (Burman et al., 2024; Van Den Berg et al., 2023).

Focusing on the interaction between phages and specialized metabolite production, we observed red colouration of the *S. albidoflavus* (formerly *S. coelicolor)* lawn surrounding undiluted spots for Ankus, DekoNeimoidia, and Naboo. Enhanced actinorhodin production in response to phage infection was also observed in previous studies (Hardy et al., 2020; Kronheim et al., 2023). However, a direct antiphage function of actinorhodin has not been reported so far. Considering the manifold triggers leading to actinorhodin production, it is currently more likely to assume that this molecule plays an important role in global stress responses and danger signalling in *Streptomyces*. For molecules of the classes of anthracyclines and aminoglycoside antibiotics – also produced by *Streptomyces* - pronounced antiphage properties have recently been described for a broad range of dsDNA phages (Kever et al., 2022; Kronheim et al., 2018). To further investigate the underlying pathways and determinants, phages able to infect naturally producing strains resistant to the compounds are needed (Hardy et al., 2023). Consequently, the reported phages add to the portfolio of phages available to understand the complex multicellular antiviral immunity in *Streptomyces*.

## Author Contributions

Conceptualization: S.E., T.L., E.D., J.F.; Data Curation, S.E.; Formal Analysis: S.E., T.L.; Funding acquisition: B.A., J.F.; Investigation: S.E., T.L.; Methodology: All; Project administration: E.D., J.F.; Resources: J.F.; Software: S.E., T.L.; Supervision: E.D., J.F.; Validation: All; Visualization: S.E., T.L.; Writing— original draft: S.E., T.L., L.K.; Writing—review and editing: All. All authors have read and agreed to the published version of the manuscript.

## Acknowledgements

The authors gratefully acknowledge the electron microscopy training, imaging and access time granted by the life science EM facility of the Ernst-Ruska Centre at Forschungszentrum Jülich. We thank Anna-Frederike Schulte-Huermann as well as Anne Fuchs-Döll, Angela Ertz, René Nork and Marcel Weckbecker from the JuLab team at the Forschungszentrum Jülich. Special thanks go to the Gesamtschule Niederzier/Merzenich student class (Luca Frinken, Anna Hasenclever, Sophie Jakobs, Anastasia Kutscherenko, Bianca Pohlen, Tim Schiffers, Jolin Schlesinger, Laurin Teschers, Lea Tirtey, Lucas Wolter, Justus Wrona and teacher Jan Schillings) for the interest and effort in sampling and isolating bacteriophages during this outreach program.

