## Supplementary Information for "Expanding the phage galaxy: Isolation and characterization of five novel *Streptomyces* siphoviruses Ankus, Byblos, DekoNeimoidia, Mandalore, and Naboo"

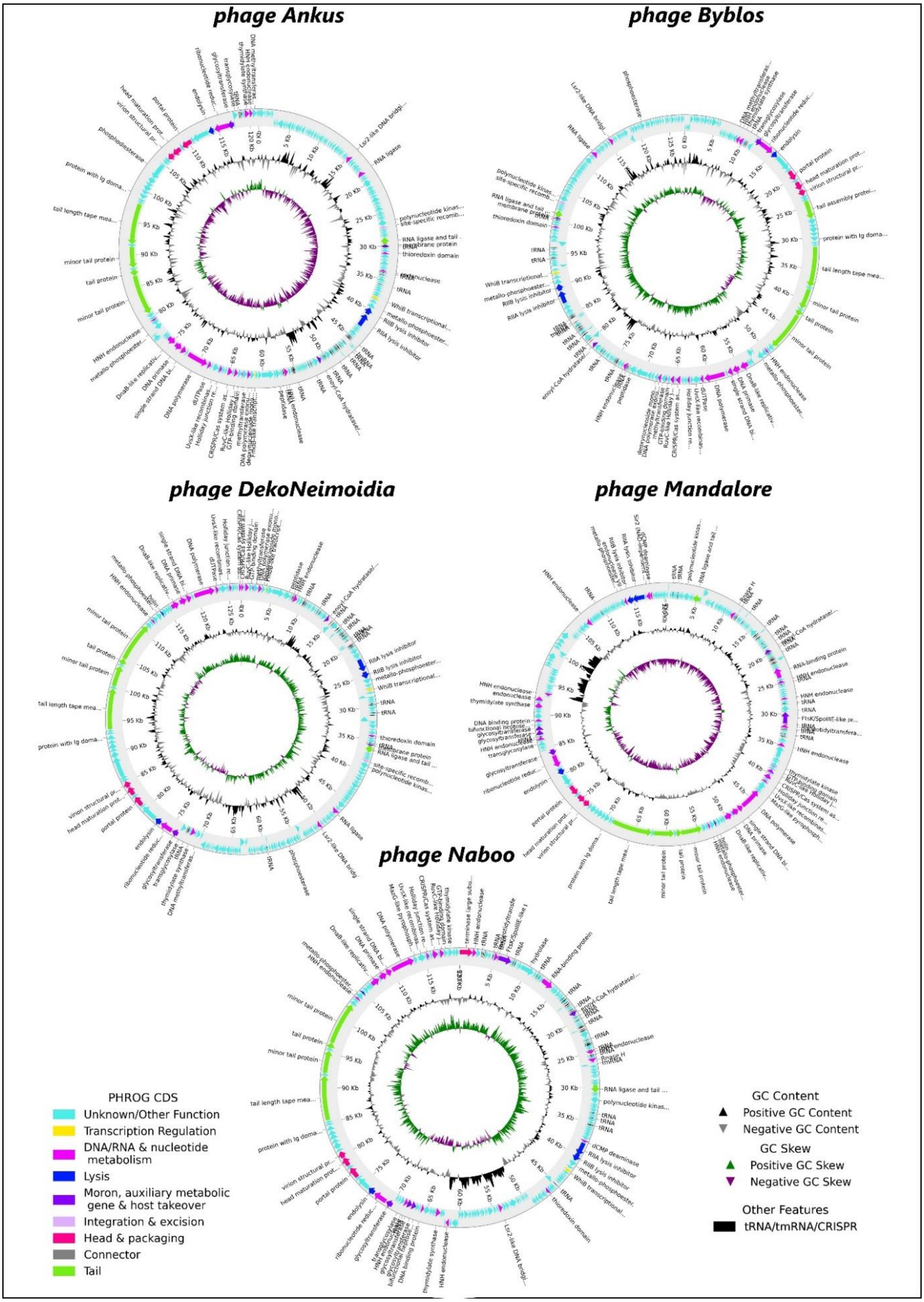


**Supplementary Figure S1: Basic genomic features of the five novel phages.** The phage open reading frames (ORFs) were predicted with Pharokka v 1.3.2 (Bouras et al., 2023) in terminase reorientation mode using PHANOTATE (McNair et al., 2019). Functional annotation was generated by matching each CDS to the PHROGs (Terzian et al., 2021), VFDB (Chen, 2004) and CARD (Alcock et al., 2019) databases using MMseqs2 (Steinegger and Söding, 2017) and PyHMMER (Larralde and Zeller, 2023). Contigs were matched to their closest hit in the

INPHARED database (Cook et al., 2021) using mash (Ondov et al., 2016). Plots were created with the pyCirclizen package.

**Supplementary Table S1: Phage particle size.** Measurements of virion particles analysed by transmission electron microscopy (TEM).

| Phage Name | Capsid diameter | Tail length | Phage size |
|---|---|---|---|
| **Ankus** | 81 nm ± 6 | 349 nm ± 8 | 430 nm |
| **Byblos** | 79 nm ± 10 | 359 nm ± 12 | 438 nm |
| **DekoNeimoidea** | 66 nm ± 3 | 358 nm ± 30 | 424 nm |
| **Mandalore** | 71 nm ± 4 | 346 nm ± 10 | 417 nm |
| **Naboo** | 75 nm ± 3 | 343 nm ± 17 | 418 nm |
